# High-throughput super-resolution analysis of influenza virus pleomorphism reveals insights into viral spatial organization

**DOI:** 10.1101/2021.09.23.461536

**Authors:** Andrew McMahon, Rebecca Andrews, Sohail V. Ghani, Thorben Cordes, Achillefs N. Kapanidis, Nicole C. Robb

## Abstract

Many viruses form highly pleomorphic particles; in influenza, these particles range from spheres of ~ 100 nm in diameter to filaments of several microns in length. Virion structure is of interest, not only in the context of virus assembly, but also because pleomorphic variations may correlate with infectivity and pathogenicity. We have used fluorescence super-resolution microscopy combined with a rapid automated analysis pipeline to image many thousands of individual influenza virions, gaining information on their size, morphology and the distribution of membrane-embedded and internal proteins. We observed broad phenotypic variability in filament size, and Fourier transform analysis of super resolution images demonstrated no generalized common spatial frequency patterning of HA or NA on the virion surface, suggesting a model of virus particle assembly where the release of progeny filaments from cells occurs in a stochastic way. Finally, we showed that in long filaments, viral RNP complexes are located preferentially within Archetti bodies, suggesting that these structures may play a role in virus transmission. Our approach therefore offers exciting new insights into influenza virus morphology and represents a powerful technique that is easily extendable to the study of pleomorphism in other pathogenic viruses.

## Introduction

Viral disease results in significant illness and deaths in humans each year, and thus represents a large healthcare and economic burden to countries around the world. The ongoing COVID-19 pandemic has resulted in the deaths of millions of people in the last year alone, while annual influenza epidemics can result in up to 650,000 respiratory deaths per year^1^, with significantly more during intermittent influenza pandemics. Despite the high mortality and morbidity associated with viruses, many aspects of their structure and morphology are poorly understood, in part due to their pleomorphic nature and small size, which represent a challenge when it comes to studying them via conventional means.

Influenza virus particles are highly pleomorphic^2^ – ranging in size from spherical virions ~ 100 nm in diameter^3^ to filaments of a similar width but reaching many micrometers in length. Similar pleomorphism has been seen for many other viruses, such as Ebola^4^, measles^5^, human parainfluenza virus type 2 (HPIV2)^6^ and Newcastle Disease Virus^7^. Filamentous strains of influenza have frequently been overlooked in virus research, both because laboratory-passaged viruses tend to be spherical (with a filamentous morphology more typical of clinical isolates) and also because ultracentrifugation and other purification and storage procedures tend to damage filaments^8^. Filamentous virus structure is of interest as it has been linked to increased pathogenicity^9–11^, resistance to neutralizing antibodies^12^ and penetration through host mucus barriers^13^, as well as offering insights into virus assembly and mechanisms of infection^14^.

Previous work has divided influenza virus particles into three broad classes; i) spherical virions with a mean outer diameter of 120 nm and an axial ratio of less than 1.2, ii) bacilliform virions of intermediate length (120-250 nm diameter), which often appear to be ellipsoidal, capsular or kidney-bean shaped, and iii) elongated filaments of greater than 250 nm in length (reviewed in^14^). Virus particles are surrounded by a lipid bilayer, in which two surface glycoproteins are embedded, the hemagglutinin (HA) and neuraminidase (NA)^15^. Electron micrographs of spherical viruses have suggested that there are ~375 surface protein spikes, of which approximately one seventh are NA and the rest are HA^15,16^. The distributions of the HA and NA glycoproteins over the virion surface are not entirely random; with studies showing that small clusters of NA are formed within the more abundant HA^15^, that NA has a tendency to cluster at the pole at the opposite end of the virion to the viral genome^15,17–19^ and that HA and NA proteins may alternate on the filament surface^13^. Beneath the lipid bilayer lies the structural matrix protein M1 - in filaments, this has been observed to have a helical structure^19^ and has been implicated as a major determinant of the pleiomorphic structure of influenza^20^. Inside the M1 core lies the viral genome, which is comprised of 8 viral RNA segments, each of which is bound by multiple nucleoproteins (NP) and the RNA polymerase (formed of the subunits PB1, PB2 and PA) to make ribonucleoprotein (RNP) complexes. Imaging work has suggested that RNP complexes cluster at one end of filamentous virus particles^13,17–19,21^. Some filamentous virions have been observed to have large bulges at one end, known as Archetti bodies^22^; previous electron microscopy images have indicated that these bulges do not appear to contain RNPs^20^ and their exact purpose in viral replication remains unknown.

The imaging of viruses by conventional fluorescence microscopy is limited as virus particles are generally smaller than the ~ 250 nm resolution limit of optical microscopy for visible light, making it impossible to gain detailed information on the shape, size and protein organisation of individual virions. Electron microscopy has represented the method of choice for high-resolution imaging of virus structure since its introduction in the 1930s, but is comparatively low throughput, time consuming and offers limited molecular identification. To overcome those limitations and facilitate the high-throughput acquisition of images of filamentous influenza virions, we used direct stochastic optical reconstruction microscopy (dSTORM)^23^, a technique that allows the location of molecules to be determined with nanometer-scale precision at a resolution of less than 20 nm and offers an exciting alternative for virus imaging^24,25^. This method is compatible with immunostaining, allowing us to specifically label viral proteins within virions.

In this study, we have combined dSTORM imaging methodology with rapid automated analysis software to carry out a high throughput and high-resolution analysis of thousands of virions at a time. We have used these tools to investigate the structure and organization of influenza H3N2 A/Udorn/72; a strain which exhibits both spherical and filamentous morphology. We found that length analysis provided a useful way of characterizing virions: filaments longer than ~ 230 nm formed a broad size distribution, while smaller particles formed two distinct populations, likely corresponding to spherical and ellipsoidal virions. In contrast to previous analyses, axial ratio analysis did not reveal distinctive populations. We also investigated the arrangement of viral proteins; demonstrating that no generalized spatial frequency patterning of HA or NA on the virion surface occurs, and observed that RNPs are preferentially located within Archetti bodies at filament ends, when Archetti bodies are present. Our analysis pipeline is versatile and can be adapted for use on multiple other pathogens, as demonstrated by its application to SARS-CoV-2. The ability to gain nanoscale structural information from many thousands of viruses in just a single experiment is valuable for the study of virus assembly mechanisms, host cell interactions and viral immunology, and should be able to contribute to the development of viral vaccines, anti-viral strategies and diagnostics.

## Results

### High throughput imaging of influenza using super-resolution microscopy

In order to establish a rapid, robust and high throughput method of imaging virus particles we focused on influenza A/Udorn/72, an influenza strain with a well-characterized spherical and filamentous phenotype^20^. Initially, we immobilised virus particles for imaging by non-specifically biotinylating the virus surface and incubating particles on the surface of a pegylated glass slide, however this method did not result in many immobilized particles (Sup. Fig. 1A), was not consistent, and required time-consuming slide preparation. To address these issues, we investigated virus immobilisation via coating the glass with the positively-charged linear polymers poly-L-lysine (Sup. Fig. 1B) or chitosan (Sup. Fig. 1C). Both chitosan and poly-L-lysine immobilised virus particles well and with low background, however poly-L-lysine was chosen for subsequent experiments due to its long-term stability and ease of preparation. Virus samples were dried directly onto glass coverslips pre-coated with poly-L-lysine, a process which took approximately 10 minutes (Fig. 1A). We showed that drying of the virus did not visibly affect filament appearance and that a combination of drying the sample and using poly-L-lysine increased the number of immobilized viruses on the slide surface (Sup. Fig. 2). Immobilised viruses were fixed, permeabilised and stained with antibodies using a standard immunofluorescence protocol (see methods).

**Fig. 1.**
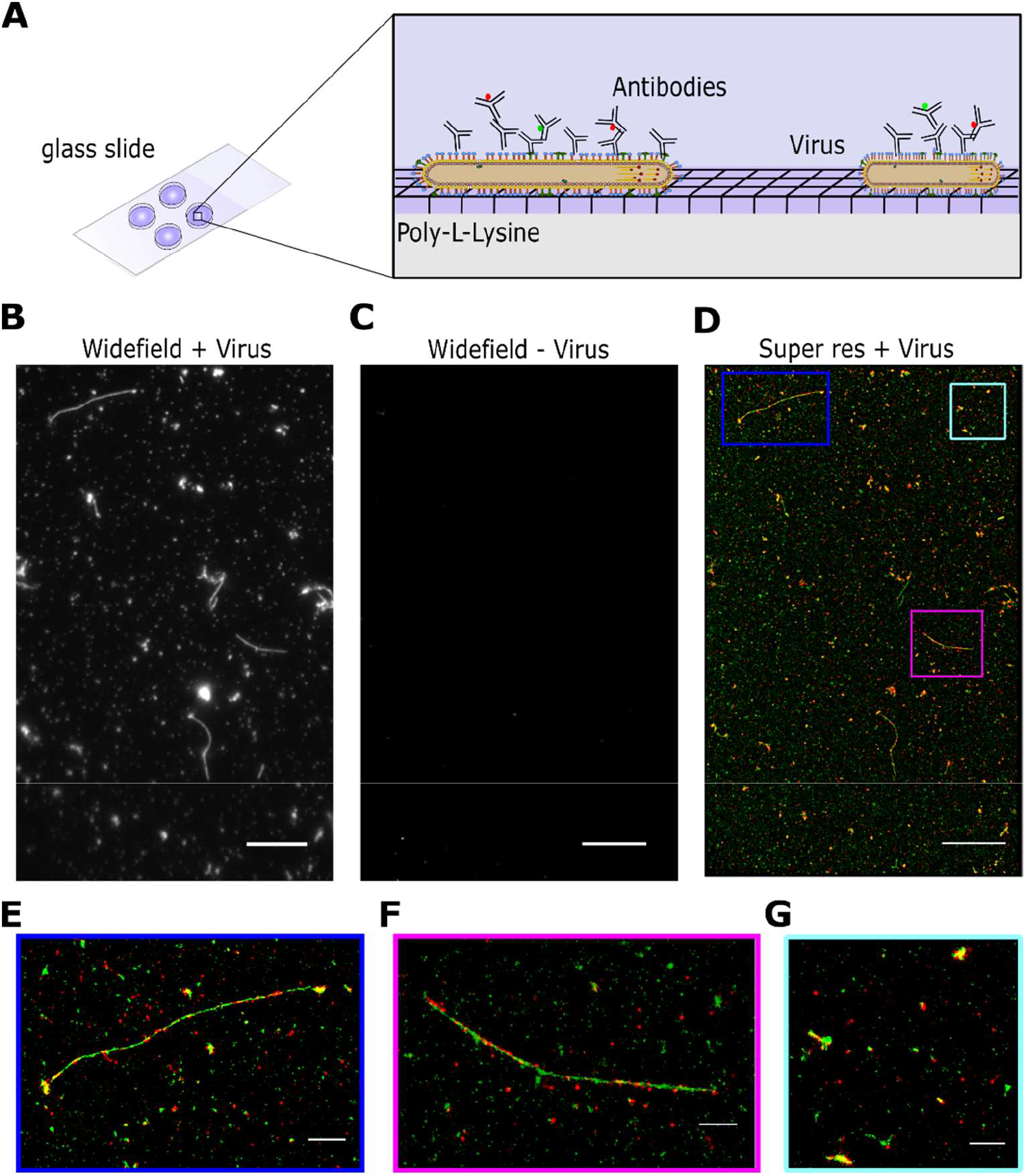
High throughput imaging of influenza using super-resolution microscopy. A) Schematic of the labelling protocol. Virus samples were dried directly onto glass coverslips pre-coated with poly-L-lysine before being fixed, permeabilised and stained with antibodies using a standard immunofluorescence protocol. B) A representative field of view (FOV) of a widefield image of labelled A/Udorn/72 influenza, imaged in the green channel. Scale bar 10 μm. C) A representative FOV of a widefield image of a virus negative sample, imaged in the green channel. Scale bar 10 μm. D) The corresponding dSTORM image of the FOV in B), where HA is labelled in green and NA is labelled in red. Scale bar 10 μm. E-G) Zoomed in images from D) showing individual filaments and spherical particles. Scale bar 5 μm.

Next, we imaged the immobilized virus particles using a widefield total internal reflection fluorescence (TIRF) microscope. A/Udorn/72 virions were dual-stained using anti-HA and anti-NA primary antibodies and secondary antibodies labelled with Alexa546 (green) and Alexa647 (red) respectively (Fig. 1B). A single field-of-view (FOV; measuring 50 x 80 μm) showed examples of multiple viruses, including large numbers of spherical particles and filaments of a wide range of different lengths (Fig. 1B). A negative control consisting of media from non-infected cells showed that the labelling was specific (Fig. 1C). A long acquisition (10,000 frames) of the FOV was taken to generate a super resolution image, which showed the virus particles at high resolution, revealing the spatial organization of HA and NA on the surface of both spherical particles and filaments (Fig. 1D-F). Multiple FOVs of the same sample can be imaged in this way, providing high-resolution information on thousands of spherical influenza virions and hundreds of filaments in each run. Taken together, this approach allows us to efficiently, rapidly and easily immobilize and image large numbers of virus particles at high resolution, in order to gain structural information on large virus populations.

### Length analysis of long filaments using diffraction-limited microscopy images

Our initial images showed a stochastic population of elongated virus particles with large variability in length, ranging from approximately 250 nm to several microns. Due to the large size of the viral filaments (larger than the diffraction limit), we initially performed a high-throughput length analysis of multiple filaments using widefield images containing diffraction-limited signals. Multiple images of A/Udorn/72 virus labelled with an anti-HA antibody were taken and a rapid automated analysis pipeline was used to measure filament length (Fig. 2A). We adjusted and binarized each image to pick out the virus filaments, using an arbitrary lower threshold of 234 nm to exclude any particles that would be below the diffraction limit. Morphological closing operations were used to fully connect the virus particles if there were regions without labelled protein and then a skeletonization operation^26^ which reduced the particles to the simplest shape to give the shape of the filaments was applied. The lengths of the resulting skeletons were taken as the lengths of the virus filaments^27,28^.

**Fig. 2.**
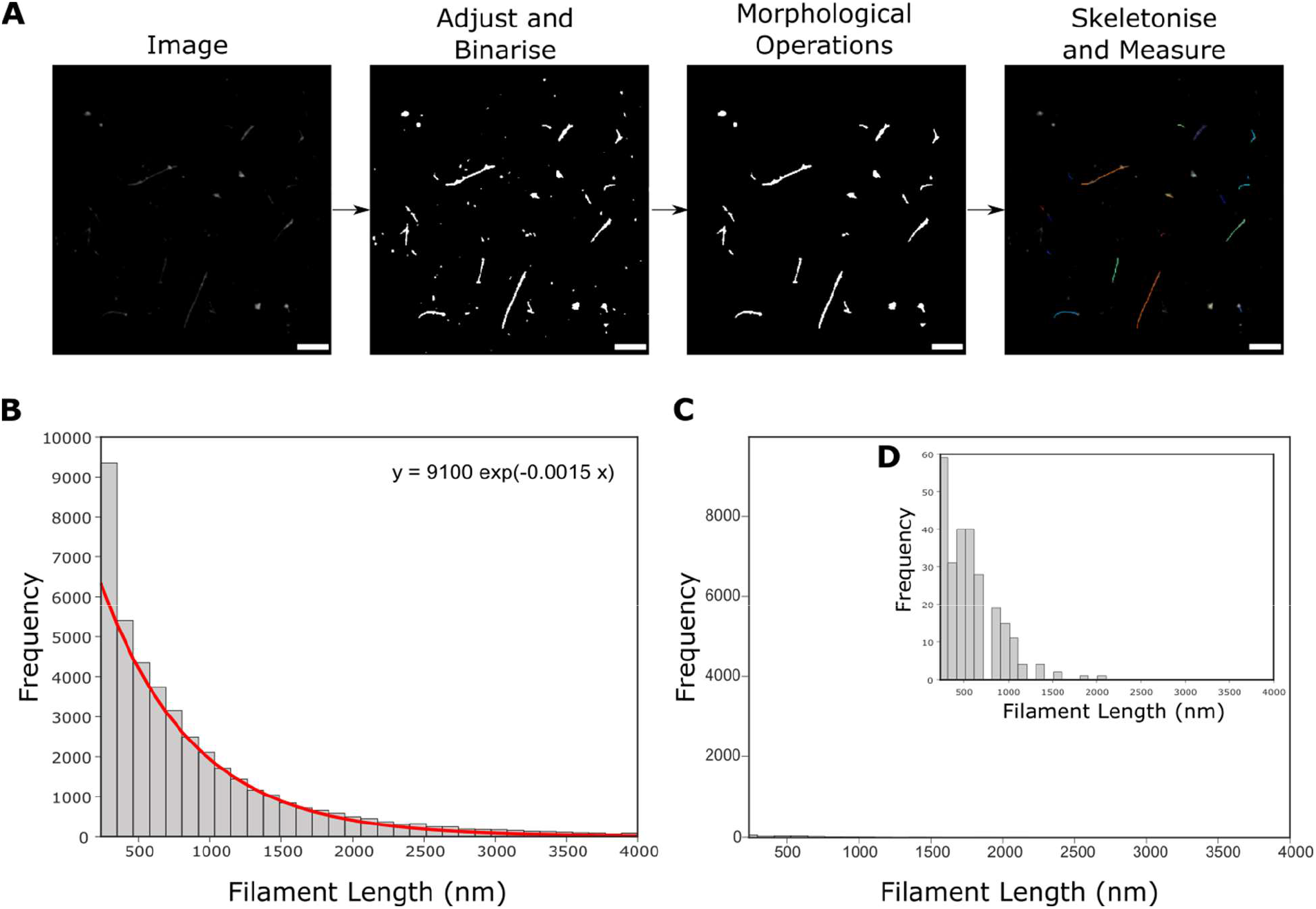
Widefield imaging and automated length analysis of A/Udorn/72 reveals that filaments fit a single distribution. A) A pipeline for analyzing filamentous Udorn particles uses multiple widefield images, which are adjusted and binarised before being skeletonized to get a measure of filament length. Scale bar 5 μm. B) Widefield images of A/Udorn/72 virus stained with an antibody against the HA protein were analysed using the pipeline in A). The resulting frequency distribution of the lengths of 46,872 filamentous particles, fit with a double exponential (red line; equation above plot). C) Histogram from the analysis of 243 virus-negative FOVs demonstrates that the background was negligible. D) Zoomed in histogram of C).

This process was completed on 486 individual FOVs, allowing the lengths of 46,872 filamentous particles to be measured. The resulting histogram was fitted with a single exponential function (Fig. 2B). Repeating the process on 243 FOVs negative for virus showed that any background signal due to spurious noise or background labelling was negligible (Fig. 2C&D). The length distribution revealed that there are significantly more filaments of shorter lengths (<1000 nm), with the frequency of filaments decreasing at longer length scales. This is consistent with previous observations that filaments are fragile^8^ and can break or fracture at longer lengths; alternatively, length may be limited by the greater amount of membrane and viral proteins required to form each filament compared to a spherical particle. The large variability in filament lengths and lack of distinctive sub-populations of filamentous virions of a particular size also suggests a model of virus particle assembly where membrane scission and the release of progeny filaments from cells occurs in a stochastic way.

### Size and shape analysis of virions using dSTORM

Having completed a comprehensive length analysis of long virions we next considered how to do this for smaller virus particles (specifically, those that fall into the spherical, ellipsoidal/bacilliform and short filament (<600 nm) categories). The small size of these viruses precluded the use of widefield images to get accurate length data, and so we turned to our super-resolution images to obtain their size distribution. A/Udorn/72 virus was immobilized, stained and imaged using dSTORM as described above; the DBSCAN clustering algorithm^29^ was used to cluster the resulting localisations in each FOV, and an ellipse was fitted to each cluster (Fig. 3A). The best-fit ellipses were used to give the major and the minor axis lengths of the clusters, which were taken as a measure of virus particle size and the results were plotted as histograms.

**Fig. 3.**
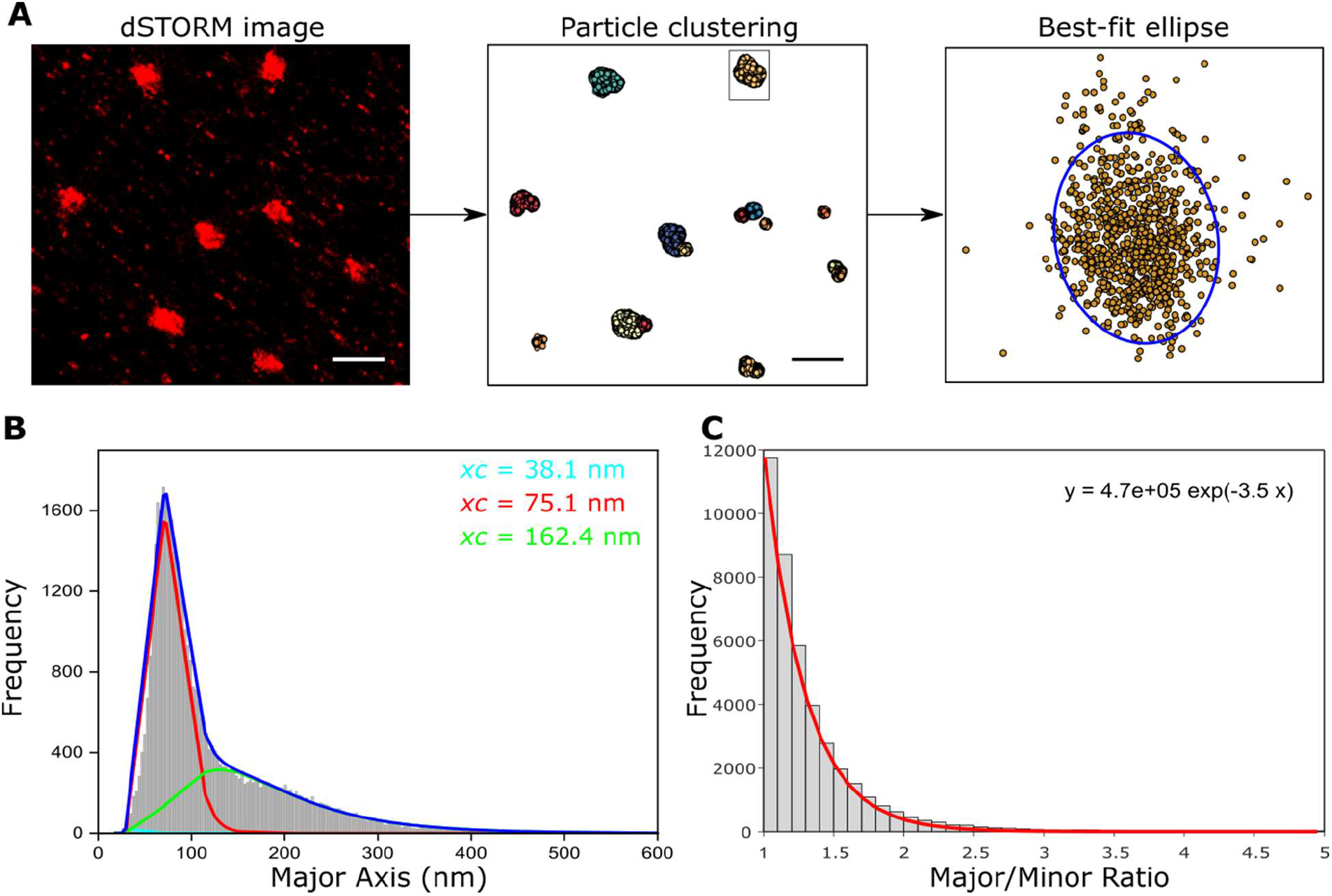
Super-resolution imaging and size analysis of spherical and bacilliform influenza particles. A) A pipeline for analyzing spherical and bacciliform influenza particles using multiple super-resolution images. Super-resolution localisations are clustered using DBSCAN and each cluster is fitted with an ellipse. Particle dimensions are extracted using the major and minor diameters of each ellipse. Scale bars 1 μm. B) A histogram of the major axis lengths shows that virions fall into two distinct populations, centered at 75.1nm and 162.4 nm. C) Histogram of the major/minor axis ratio shows a single distribution.

This process was completed on 46 individual FOVs, allowing the size of 41,754 particles to be measured. By repeating the process on virus-negative FOVs, the data was corrected for the majority of points due to noise or background (Sup. Fig. 3). The resulting histogram of the major axis length was fitted using a triple log-normal function, chosen to take into account the two distinctive virion populations as well as a minor third distribution representing the small amount of background signal still remaining (Fig. 3B). This analysis revealed two distinct populations of virions, centered at ~ 75 nm (54.6% of the total particles) and ~ 160 nm (44.2% of the total particles), likely corresponding to spherical and ellipsoidal virions respectively. The third distribution, representing 1.2% of all particles, was centered at ~ 40 nm; the extremely small size of this distribution suggests that our clustering methods and subtraction of data points from virus-negative experiments were sufficient to remove most non-specific localisations. The primary source of error in our measurement of length scales is due to the localization error from the Gaussian fit to each localisation during super-resolution image reconstruction. We therefore plotted the error in individual localisations for the super-resolution images used in this analysis, which gave a median error of 7.4 nm in the x and 7.2 nm in the y direction, too small to make any meaningful difference to our results (Sup. Fig. 4).

EM images of virus particles have previously been categorized based on their axial ratio (< 1.2 for spherical virions, > 1.2 for bacilliform virions and filaments), as well as length. We therefore plotted the major/minor axis histogram from the measured particles, and fitted the resulting plot with a single exponential function (Fig. 3C). Rather than being able to distinguish two distinct axial ratio populations (of > and < than 1.2) we observed a single population. This suggests that when analysing large numbers of virions (many thousands compared to a few hundred via EM) a much larger variability in particle shape may be captured, thus suggesting that a greater heterogeneity in virus shape than previously thought may occur.

In order to demonstrate the versatility of our imaging and analysis pipelines, we carried out similar experiments on the SARS-CoV-2 virus, the causative agent of the Covid-19 pandemic. Fixed SARS-CoV-2 virions were dual-labelled with anti-spike and anti-nucleoprotein (N) primary antibodies and secondary antibodies labelled with Alexa647 (red) and Alexa546 (green) respectively (Fig. 4A). Multiple co-localised particles were observed (Fig. 4B), corresponding to double-labelled virions, as well as particles labelled in just the green channel (Fig. 4C), corresponding to virions labelled for nucleoprotein but lacking spike protein, likely lost from the surface during sample preparation. N protein localisations were chosen to give an estimate of virion size as particles have been shown to have only small numbers of spike proteins on the surface (approximately one per 1,000 nm^2^ of membrane surface, compared with approximately one per 100 nm^2^ for influenza^30^). Super-resolution localisations in the green channel (labelling the N protein) were therefore clustered and each cluster fitted with an ellipse to extract particle dimensions. A histogram of the major axis lengths fitted with a Gaussian function shows that virions fall into a single population centred at ~ 144 nm (Fig. 4D), in keeping with an expected size of SARS-CoV-2 virus particles of 50 – 200 nm in diameter^31^, while the major/minor axis histogram from the labelled SARS-CoV-2 particles showed a tighter distribution of virion axial ratio than that found for influenza, in keeping with observations that SARS-CoV-2 forms largely spherical particles (Fig. 4E). These results confirm previous findings on the dimensions of the SARS-CoV-2 virus and demonstrate the general applicability of our analysis pipeline to multiple other viruses.

**Fig. 4.**
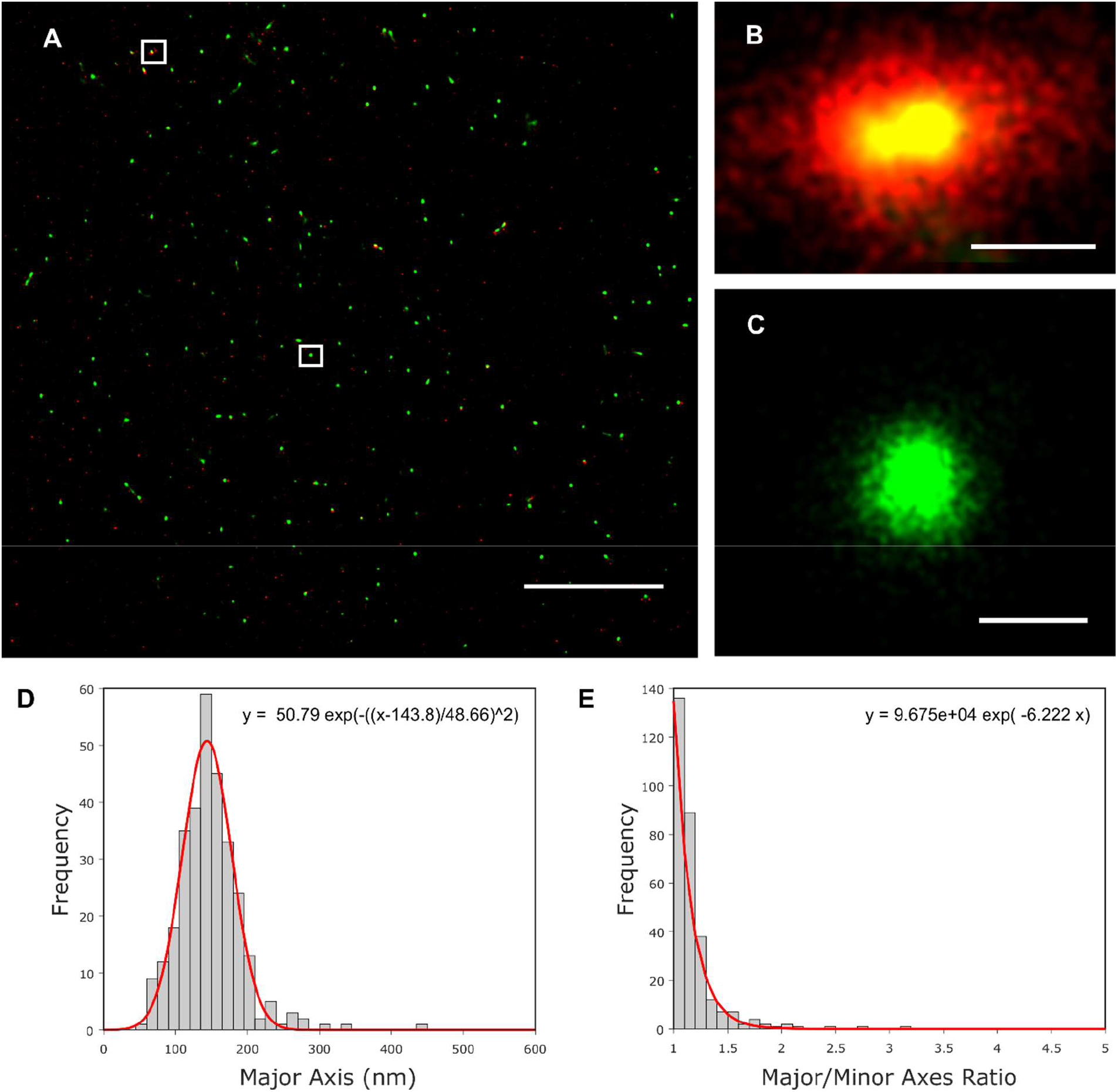
Super-resolution imaging and size analysis of SARS-CoV-2 virus particles. A) A representative super-resolution image of SARS-CoV-2 virions dual-labelled with anti-spike and anti-nucleoprotein (N) primary antibodies and secondary antibodies labelled with Alexa647 (red) and Alexa546 (green) respectively. Scale bar 10 μm. B&C) Zoomed images of individual SARS-CoV-2 particles (highlighted in the white boxes in A). Scale bar 100 nm. D) Super-resolution localisations in the green channel (labelling the N protein) were clustered and each cluster fitted with an ellipse to extract particle dimensions. A histogram of the major axis lengths fitted with a Gaussian function shows that virions fall into a single population, centred at 143.8nm. E) Histogram of the major/minor axis ratio shows a single distribution.

### Surface protein organization through Fourier transform analysis of super resolution images

The distributions of the HA and NA glycoproteins over the virion surface are not entirely random; indeed, when we stained filaments for either HA (Fig. 5A) or NA (Fig. 5B) the resulting super-resolution images showed a non-uniform distribution of protein signals. We therefore used intensity analysis and Fourier transforms to investigate the patterning of surface proteins on a large number of filaments. A/Udorn/72 was immobilised, labelled and imaged using dSTORM; the filaments were fit with skeletons as above and the intensity profiles of filaments were found by fitting the skeleton with a polynomial, finding the normal, summing the intensity over the normal and plotting this over the length of the filament (Fig. 5C). Finally, the overall patterning was shown by summing the Fourier transforms of all of the filaments, in order to form an average distribution of the patterning.

**Fig. 5.**
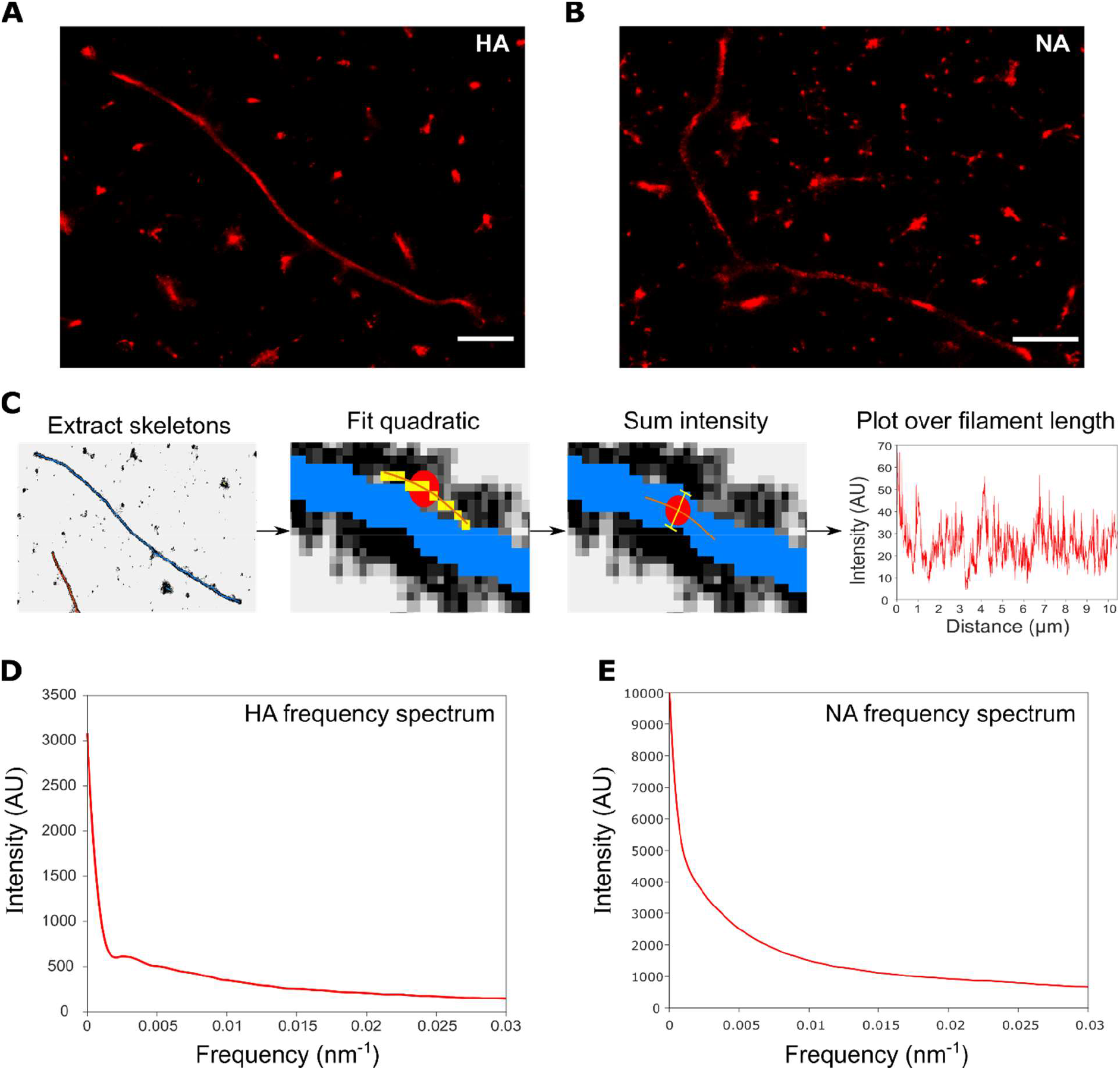
Analysis of the frequency distribution of influenza surface proteins. A&B) Representative super-resolution images of filaments labelled with A) HA antibody and B) NA antibody. Scale bars 1 μm. C) A pipeline for analyzing the influenza filament protein distribution using multiple super-resolution images. Super-resolution images are skeletonized, fit with quadratics at a point along the skeletons; the normal is found and the intensity across this normal is summed and the Fourier transform is taken and summed across filaments. D) The frequency distribution of the HA proteins from 1,067 filaments from 8 FOVs. The frequency spectrum for HA has no distinctive peaks, suggesting that there is no common spatial patterning of HA across filaments. E) The frequency spectrum for NA also has no distinctive peaks, suggesting that there is no common spatial patterning of NA across filaments.

We started by analyzing the frequency distribution of the HA protein. The resulting distribution showed no peaks apart from the high signal at close to 0 nm^-1^ which is due to the spatial frequency of the filament as a whole (Fig. 5D). This suggests that there is no set frequency at which HA proteins are located along influenza virions, which in turn suggests that the surface virion proteins are not patterned deterministically. The broad distribution at low frequencies indicates that individual filaments can have patterning at a certain spatial frequency, but it is not common among all filaments. Similar analysis was conducted for the NA protein (Fig. 5E), with the same conclusion that no generalized NA spatial patterning on influenza filaments occurs.

To confirm our observation that no spatial frequency arrangement of the surface proteins occurs, filamentous virions with a length of 500 nm and a diameter of 80 nm were simulated through Monte Carlo methods (Sup. Fig. 5). Initially, points corresponding to the HA and NA proteins were positioned randomly on a cylinder with hemispherical caps; by projecting these localisations into two dimensions and randomly fitting points with a normal distribution around the protein positions, simulated dSTORM images could be built up (Fig. 6A). For comparison, we simulated dSTORM images of filaments with a forced frequency of protein alternation, representing filaments with a deterministic protein placement (Fig. 6B). The relative positioning of the surface proteins in our simulations was confirmed by plotting the HA intensity profiles of representative simulated filaments with either randomly distributed HA and NA, or with alternating HA and NA (Fig. 6C&D). Next, we investigated the spatial patterning of these simulated filaments by computing the sum of the Fourier transforms of the histograms of protein location along 1000 randomly distributed, and 1000 alternating, simulated filaments. The frequency distribution for filaments with completely randomly chosen points (Fig. 6E) was similar to the distributions obtained from experimental data for the HA and NA proteins (Fig. 5D&E), with no peaks and a broad spread of frequencies at low numbers. The plot for simulations with a forced alternation of 0.01 nm^-1^ however, showed a peak at 0.01 nm^-1^ and a further small peak of the harmonic at 0.02nm^-1^ (Fig. 6D). Together, our data suggests that no deterministic spatial patterning of the HA and NA glycoproteins occurs over a large population of filaments.

**Fig 6.**
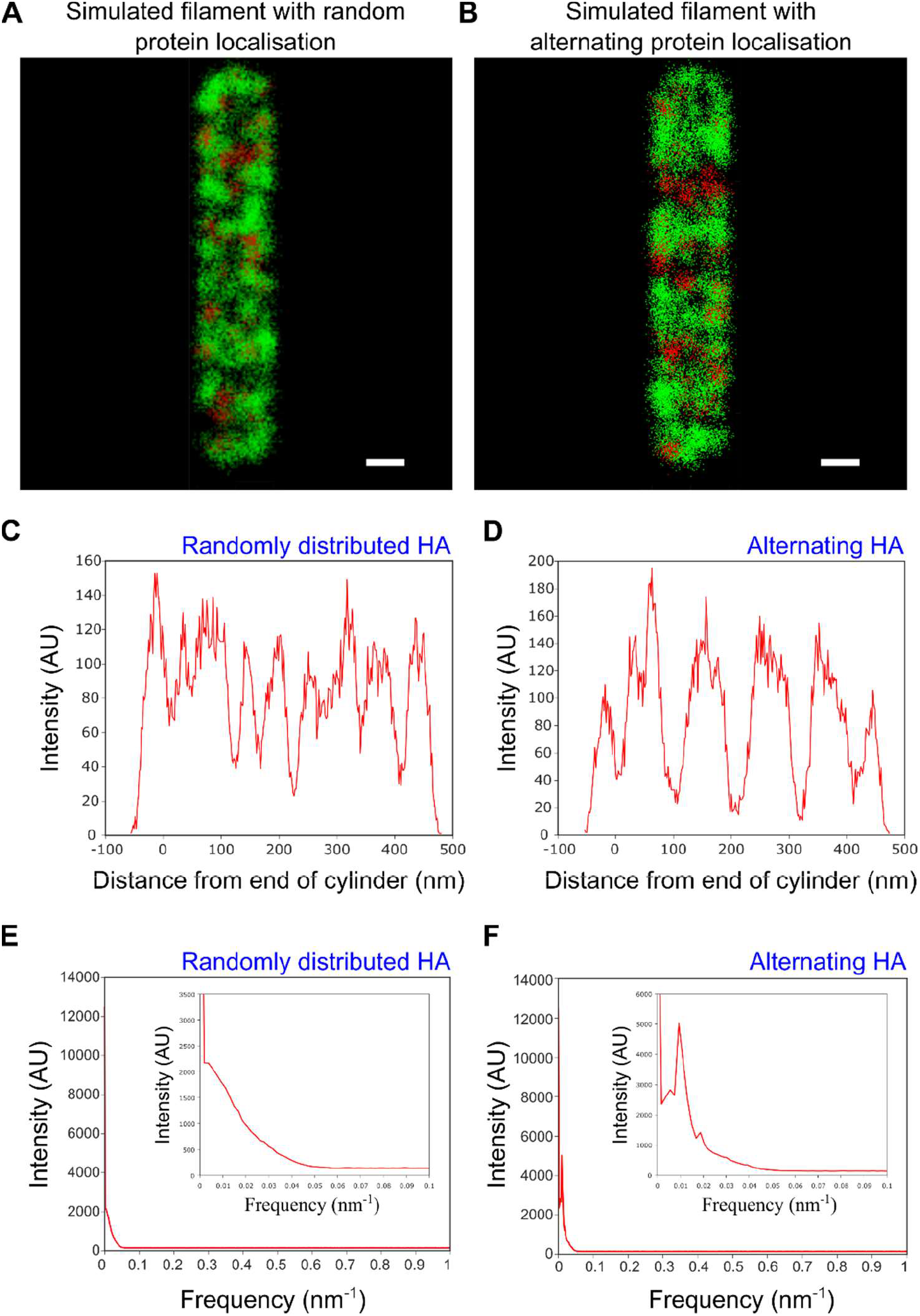
Filamentous virion simulations and analysis of alternation on simulated dSTORM images. A&B) Representative simulated dSTORM images of filamentous influenza particles with A) random protein placement and B) 0.01 nm^-1^ frequency alternating protein placement. HA is shown as green and NA as red, on a filament of 500 nm in length and 80 nm in width. Scale bars 50 nm. C) Intensity profile for a representative simulated filament with randomly distributed HA and NA. D) Same as C) but for filaments with alternating HA/NA. E) The sum of the Fourier transforms of the intensity plots of 1000 simulated filaments with randomly distributed HA and NA, suggesting no common patterning exists. F) Same as E) but for filaments with alternating HA/NA. The peak at 0.01nm^-1^ and harmonic at 0.02nm^-1^ confirm that protein patterning can be detected using this method.

### Single-molecule FISH reveals single RNP complexes at the ends of filaments

Electron microscopy images of bacilliforms and short filaments of influenza have suggested that the RNPs that contain the viral RNA cluster at one end of the virus interior, however clear images of the genome have only been obtained in a minority of longer filaments (reviewed in^8^). In order to clarify the location of RNPs within filaments we used single-molecule fluorescence in situ hybridization (smFISH) to specifically label the influenza NA segment RNA^32^ combined with antibody labelling against the HA protein to outline the filament shape. smFISH uses an array of fluorescently labelled DNA probes that bind to several sites on the viral RNA, thereby accumulating several fluorescent dye molecules on a viral RNA, making it visible as a bright spot.

A/Udorn/72 virus was immobilized and stained for HA (green) before being incubated overnight with Quasar 670 (red)-labelled FISH probes. Widefield images of the sample showed multiple long filaments of up to several microns in length (green), which were then overlaid with super-resolution localisations from the FISH probes against the NA gene segment (red) (Fig. 7A&B). 83 filaments were imaged in total, of which 15 were manually discarded as they overlapped with other filaments, making analysis challenging. Out of the remaining 68 filaments, 14 (20.1%) of these showed a distinctive bulbous shape at one end in the green channel, corresponding to Archetti bodies (Fig. 7A; white arrows), while the remaining filaments had no visible bulge at the end (Fig. 7B).

**Fig. 7.**
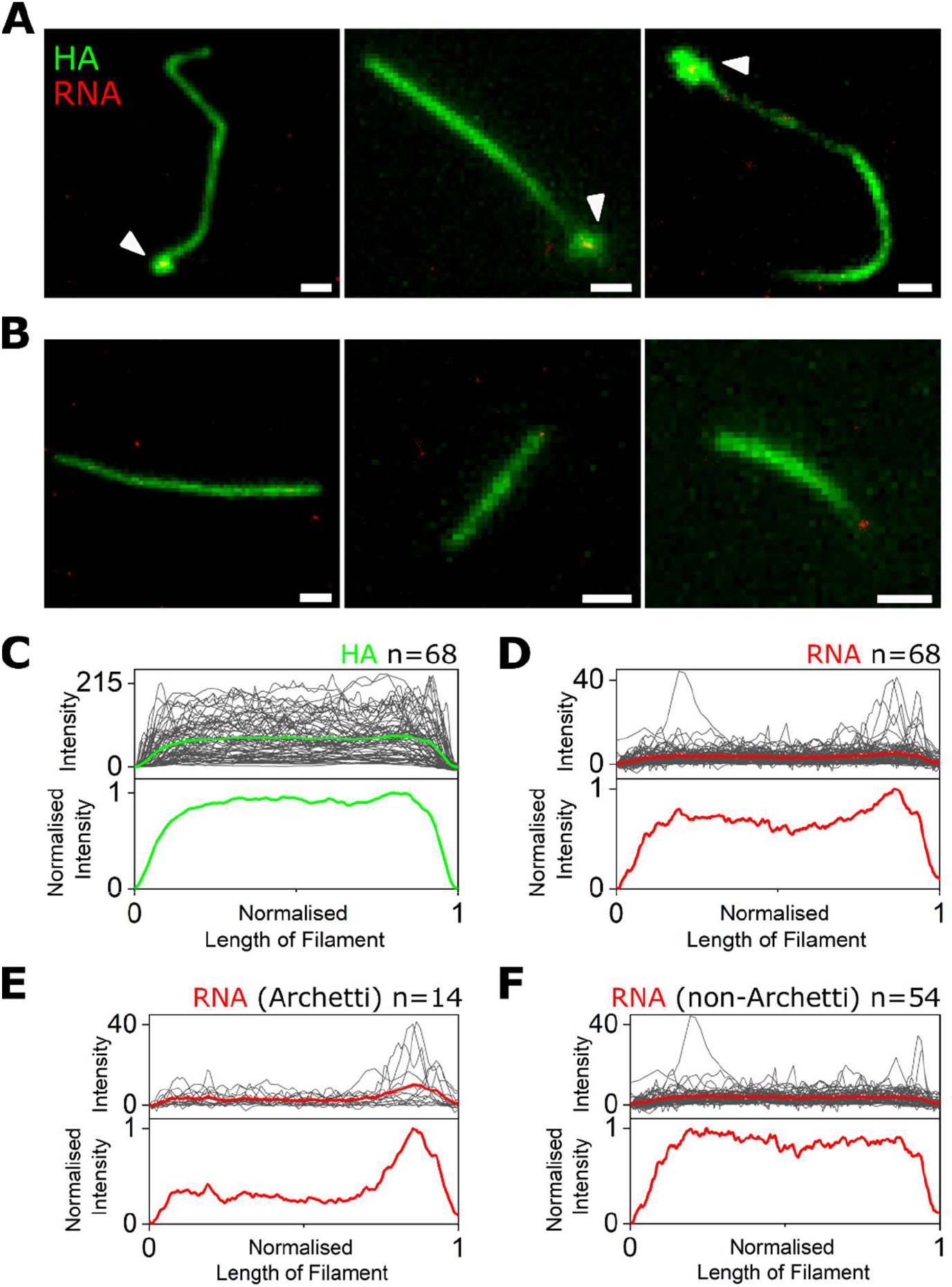
RNA is located at one end of influenza filaments. A&B) Diffraction limited images of filamentous A/Udorn/72 stained with an anti-HA antibody (green) overlaid with super-resolution localisations from an array of FISH probes against the NA gene segment (red). Scale bars 1 μm. White arrows denote Archetti bodies. C) Top: Intensity traces (grey) of the HA signal from 68 filaments, with the average intensity profile shown as a green line. Bottom: Normalised average HA signal from 68 filaments. D) Same as C) but for the red super-resolution signal corresponding to NA RNA from the 68 filaments. E) Top: Intensity traces (grey) of the RNA signal from 14 filaments with a visible Archetti body, with the average intensity profile shown as a green line. Bottom: Normalised average RNA signal from the 14 filaments. F) Same as E) but for the RNA signal from 54 filaments with no visible Archetti body.

In order to determine the position of the RNA in each filament, a line was drawn along the axis of the filaments (using the green HA signal to approximate the filament outline) and the intensity profiles of the line in both the green and red channels was measured. The averaged intensity of the HA signal from all 68 filaments analysed was uniformly distributed over the length of the filaments (Fig. 7C), however the averaged intensity traces of the super-resolution signal corresponding to NA RNA from the 68 filaments showed distinct peaks with a tendency to be located towards the filament ends (Fig. 7D). This trend was more obvious for filaments that had Archetti bodies (Fig. 7E and Sup. Fig. 6A), compared to those without (Fig. 7F and Sup. Fig. 6B); and all filaments with Archetti bodies had a FISH signal corresponding to viral RNA within the structure. This analysis suggests that Archetti bodies may play a role in housing RNPs and potentially in virus transmission.

## Discussion

In this work, we have shown that a combination of super-resolution imaging and high-throughput analysis of virus particles allows for the structural features of multiple virions to be examined at a time. Using the highly pleomorphic influenza strain A/Udorn/72 as a proof of principle, we imaged thousands of filamentous and spherical virions, allowing the features of a large population of virus particles to be ascertained at high-resolution. This approach offers several advantages over electron microscopy or conventional fluorescence microscopy, which can be low through-put or lack the resolution required to resolve virus particle features. We verified the utility of our analysis pipeline for multiple virus types using the well-characterised SARS-CoV-2 virus, which forms laregly spherical virions of ranging from 50 – 200 nm in size^31^.

Large viral filaments that measured over 250nm in length were analysed using widefield microscopy; analysis of the size distribution of >40,000 filamentous virus in this way showed a broad range of filament sizes. The frequency of filaments decreased with length, suggesting that long filaments are easily broken or that production of longer filaments may be restricted by the need for the large amounts of viral protein required to form each filament compared to a spherical particle. Virion size at short length scales was investigated using super resolution microscopy, where the analysis revealed two distinct size distributions, in keeping with previous observations that influenza forms both spherical and bacilliform particles^7^. We weren’t able to distinguish between these two populations using their axial ratio, instead observing just a single population of viruses, perhaps because our analysis of such a large number of virions captured a greater heterogeneity in particle shape than previously observed.

Our analysis demonstrated large variability in filament length, as well as a lack of distinctly sized subpopulations within a filamentous virus population. In influenza, the initiation of virus budding is thought to be initiated by clustering of HA and NA in lipid raft domains on the cell surface, followed by recruitment of M1 which serves as a docking site for the RNPs just below the plasma membrane^33^. Elongation of the bidding virion is caused by polymerization of the M1 protein, and the viral M2 protein is thought to localize at the periphery of the budding virus through interactions with M1^33^. Finally, membrane scission, mediated by M2, leads to release of the budding virus^34^. The regulation of filament formation is not fully understood, but appears to be a complex process driven by at least M1 and M2, and possibly other viral or cellular proteins. Our analysis suggests a model of virus particle assembly where the release of progeny filaments from cells occurs in a stochastic way, leading to filaments of a wide range of lengths. This model is supported by a previous observation that influenza produces virions with broad variations in size and protein composition, which led to the suggestion that phenotypic variability may contribute to virus survival under stress conditions such as in the presence of antiviral NA inhibitor drugs^35^.

Electron microscopy images of influenza virions allow NA and HA spikes on the viral surface to be distinguished by length and density, consistent with their known crystallographic structures, leading to the observation that HA and NA separate into distinct clusters. In addition, NA has been shown to preferentially cluster at the filament pole, often at the opposite end to the RNPs^13,19^, and super-resolution analysis of labelled HA and NA has suggested possible alternation of the two proteins^13^. Our analysis shows that there isn’t a common spatial frequency patterning observed amongst filaments, suggesting that HA/NA patterning isn’t driven deterministically. We also showed that in long filaments, viral RNP complexes are located preferentially towards the filament pole within Archetti bodies, when Archetti bodies are present. Further work on these structures may lend support to the hypothesis that Archetti bodies may play a role in virus transmission.

In this work, we have shown that high-resolution microscopy and rapid automated analysis software can be used to gain structural information on thousands of virions at a time. Filaments can cause difficulties in filtering and purifying influenza viruses during vaccine purification, suggesting that the ability to rapidly assess and quantify virus size and morphology may be useful in vaccine production. Our methods may also be useful for future studies of virus assembly and budding, cell-to-cell transmission and virus pathogenesis, or the development of viral diagnostic strategies that rely on virus morphology for identification.

## Methods

### Virus strains

The influenza strain H3N2 A/Udorn/72 (Udorn) was grown in Madin-Darby Canine Kidney (MDCK) cells as described previously^36^. To produce filament-containing stocks for analysis, confluent MDCK cells were infected at a multiplicity of infection of 0.001 and incubated at 37°C in serum-free media (Dulbecco’s Modified Eagle Medium, Gibco) supplemented with 2 μg/mL TPCK-treated trypsin (Sigma) for 24 hours. Supernatants were harvested and clarified at 4000 rpm for 5 minutes at room temperature to remove cell debris before being used without any further purification or concentration steps. 0.2% formaldehyde was added and the supernatant was stored at 4°C to prevent filament degradation. SARS-CoV-2 was grown in Vero E6 cells and collected as above. The virus was inactivated by addition of 4% formaldehyde before use^37^ and stored at −80°C.

### Virus immobilization

Unless otherwise specified, viruses were immobilized using poly-L-lysine. Coverslips (25×65mm, thickness number 1, VWR) were heated in a furnace at 500°C for 1 hour to burn off any dust. A silicone gasket (Grace Bio-Labs, USA) was added to the glass slide and 0.01% poly-L-Lysine (Sigma) solution was added to the wells for 30 minutes. The excess was removed and the chambers were washed three times with MilliQ water and the slide allowed to dry. As an alternative chitosan was used in place of poly-L-lysine (0.015% chitosan powder (Sigma) in 0.1M ethanoic acid).

For specific immobilisation of viruses via biotinylation, passivated microscope slides were prepared by washing in acetone and Vectabond solution (Vector Laboratories) before being incubated with NHS-PEG:Biotin-NHS-PEG in an 80:1 ratio. 0.5 mg/mL neutravidin was incubated for 10 minutes at room temperature on the slide shortly before virus was added. Viruses were biotinylated by incubation in a 1 mg/mL Sulfo-NHS-LC-Biotin (ThermoFisher) for 3 hours at 37°C before being fixed and immunolabelled as described below.

### Sample preparation

Virus was incubated in the wells at 45°C until the well was dry. Immobilised virus was fixed with 4% formaldehyde (Thermo Scientific) in phosphate buffered saline (PBS) for 10 minutes at room temperature, before being permeabilised in 0.5% Triton-X-100 (MP Biomedicals) for 15 minutes. The sample was blocked to prevent non-specific binding with 10% donkey serum (Sigma) and 0.1% Tween-20 in PBS for 1 hour at room temperature. Primary antibodies were diluted in blocking buffer and incubated with the sample for 1 hour, followed by washing in PBS and labelling with secondary antibodies (Invitrogen). A final washing step was carried out and samples were stored in PBS at 4°C until imaging. For super resolution imaging, the PBS was replaced with a STORM imaging buffer comprising an enzymatic oxygen scavenging system consisting of 1 mg/mL glucose oxidase and 40 μg/mL catalase, 10% glucose and 0.05 M mercaptoethylamine (MEA) in 1x PBS. Multiple independent repeats, with different virus preparations, were taken over different days.

A/Udorn/72 virions were labelled with the mouse anti-HA primary antibody Hc83x (a kind gift from Stephen Wharton, Francis Crick Institute) and goat anti-NA and anti-M1 (a kind gift from Jeremy Rossman, University of Kent). SARS-CoV-2 virions were labelled with the human anti-spike antibody EY6A^38^ (a kind gift from Tiong Tan and Alain Townsend, University of Oxford) and a SARS-CoV-2 nucleocapsid antibody (GTX632269) from Genetex. Secondary antibodies labelled with either Alexa647 or Alexa 546 were purchased from Invitrogen.

For fluorescence in situ hybridisation, the samples were immunostained for HA as described above, then fixed again with 4% formaldehyde in PBS for 10 minutes, washed with 2x saline sodium citrate (SSC) and subsequently permeabilized with 0.5% Triton X-100 in PBS for 10 min. After washing with 2x SSC, the surface was blocked with blocking buffer (2x SSC, 10% formamide, 10% dextran sulfate, 0.02% RNase free-BSA, 0.2 mg/ml E. coli tRNA, 1% RNasin Plus) for 30 minutes at 37°C. The sample was then incubated with 1 μM FISH probes (Stellaris LGC; 48 probes designed against the NA gene segment and labelled with Quasar 670) in blocking buffer for 1-3h at 37°C. After hybridization, the sample was washed thrice with 2x SSC, 10% formamide, 1 % RNasin Plus and then washed three times with 2x SSC before being imaged.

### Imaging

All experiments were performed on a commercially available Nanoimager fluorescence microscope (Oxford Nanoimaging). The sample was imaged using total internal reflection fluorescence (TIRF) microscopy. The laser illumination was focused at an angle of 53° with respect to the default position. Images of a field of view (FOV) measuring 80 x 50 μm were taken with an exposure time of 30 ms. For super resolution imaging the laser intensities were gradually increased up to a maximum of 780 kW/cm^2^ for both the green (532 nm) and red (647 nm) lasers, and movies of between 5,000 and 15,000 frames were taken. The red channel was imaged first, followed by the green channel.

### Filament length analysis

Images were saved as .tif files, opened in ImageJ and each FOV was cropped so that just the single channel in which the HA protein was labelled was used. The images were adjusted using the MATLAB imadjust function and binarized using the MATLAB imbinarize function with a threshold of 0.001. The background of the resulting black and white image was removed using the bwpropfilt MATLAB function and the resulting image was closed, skeletonised^26^ and labelled using the MATLAB bwmorph, bwskel and bwlabel functions respectively. Any skeletons of less than 234nm (2 pixels) long were excluded from the analysis, as were overlapping skeletons (due to overlapping filaments dried onto the slide. Results were plotted as a histogram and fitted with a bi-exponential function, detailed on the graph.

### Spherical size analysis

Each FOV was drift corrected, and super-resolution localisations were extracted in each frame using the inbuilt Nanoimager software. The localisations were exported and analysed further in Matlab. Localisations were clustered using the sklearn library implementation of the DBScan clustering algorithm with a minimum cluster size of 200 and an epsilon of 30nm^29^. The clusters were fit using the confidence_ellipse library method of fitting a confidence ellipse to a set of points with 2.0 as the standard deviation of the ellipse. This algorithm uses the covariance of a set of points, the Pearson coefficient and the mean of the points to fit an ellipse to a set of points. The major and minor axes of the ellipse were taken as the size of the viral particles.

### Protein distribution and spatial frequency analysis

FOVs with multiple filaments (> 615 nm) labelled with the protein of interest (HA or NA) were skeletonized as in the filament length analysis. For each pixel in the skeleton of filaments, a fit line was produced (using the MATLAB polyfit function) using the next 5 points in each direction from the pixel of interest. From this quadratic, the normal was found and the intensity of the pixels in the normal were summed. After iterating this over the skeleton, the calculated line profile of each filament were Fourier transformed using the MATLAB fft function and the resulting distributions across all filaments were summed to produce the average frequency spectrum for that surface protein^39^.

### Virion simulation

dSTORM images of viruses were simulated using a Monte Carlo method with the probability of a protein being placed equally across the virion and an exclusion distance of 20nm. Filaments were modelled as cylinders with hemispherical caps. After projecting the simulated filaments into 2D, localisations were randomly placed with a normal distribution about the protein location with a standard deviation of 7.4nm (derived from the localization precision of our data) and the image was coloured as a STORM image^40^. Alternating proteins were placed by choosing the random number for the protein location along the filament body from probability distributions derived from the functions |sin(kx/2)| and |cos(kx/2)|.

### FISH image analysis

The inbuilt Nanoimager software was used to create dual-colour images of diffraction-limited green fluorescence and super-resolution localisations of red fluorescence of the filaments. To create the intensity along the filament figures, movies recorded at 30 ms exposure with 1000 frames were projected in FIJI to make a single summation image. In FIJI, a line was taken, with no directionality, along the axis of the filaments measuring the intensity in the green and red emission channels of the microscope. The background intensity (the first value in the intensity line) was subtracted to produce the raw intensity of the fluorescence, the length in pixels was normalised (0-1), and graphs plotted for all lines.

## Acknowledgements

Special thanks to Dr. Edward Hutchinson and Dr. Jack Hirst from the University of Glasgow for fruitful discussions. We also thank Stephen Wharton, The Francis Crick Institute, Dr. Jeremy Rossman, University of Kent and Tiong Tan, Lisa Schimanski, Pramila Rijal and Prof. Alain Townsend, University of Oxford, and Kuan-Ying A. Huang, Chang Gung Memorial Hospital, Taiwan, for their generous gifts of antibodies. Many thanks also to Dr. Rebecca Moore and Prof. William James from the Sir William Dunn School of Pathology, University of Oxford, for providing SARS-CoV-2. This research was supported by Royal Society Dorothy Hodgkin Research Fellowship DKR00620 and Research Grant for Research Fellows RGF\R1\180054 (to N.C.R.), a BBSRC-funded MIBTP PhD studentship (to S.V.G), the University of Oxford COVID-19 Research Response Fund and BBSRC grant BB/V001868/1 (to N.C.R. and A.N.K.) and Wellcome Trust grant 110164/Z/15/Z (A.N.K.). The authors gratefully acknowledge the Micron Advanced Bioimaging Unit (supported by Wellcome Strategic Awards 091911/B/10/Z and 107457/Z/15/Z) for their support & assistance in this work.

## Author Contributions

A.M. contributed experimental design, data acquisition, software, data analysis and manuscript drafting; S.V.G. contributed simulation software; R.A. contributed data acquisition and analysis; T.C. contributed to project conception; A.N.K. contributed to data interpretation and N.C.R. contributed to project conception, experimental design and data interpretation. All authors reviewed the manuscript.

## Data Availability

Data and analysis code is available at https://github.com/amcmahon1345/filaments.git.

