## Supplemental figures for "High-throughput super-resolution analysis of influenza virus pleomorphism reveals insights into viral spatial organization"

**Supplementary Figures**

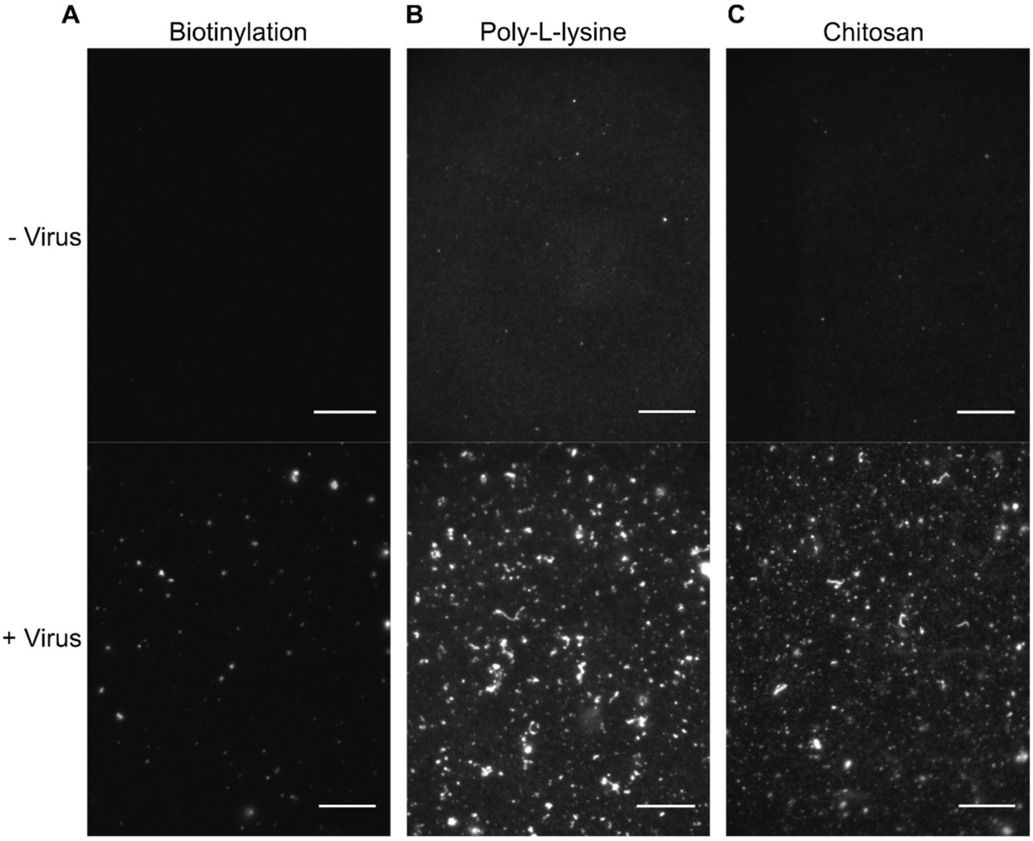


**Sup. Fig. 1. Immobilisation methods for filamentous viruses.** A) A virus negative control (top) or an A/Udorn/72 virus sample (bottom), were immobilized via a specific biotin/PEG linkage. Virus particles were biotinylated by incubation with 1 mg/mL Sulfo-NHS-LC-Biotin (ThermoFisher) for 3 hours at 37°C before being immobilised on a pegylated slide. The virus was labelled with an anti-Udorn primary antibody and an Alexa647 secondary antibody and imaged on a widefield TIRF microscope. Scale bars 10μm. B) A virus sample was incubated on a slide pre-treated with 0.01% poly-L-lysine. C) A virus sample was incubated on a slide pre-treated with 0.015 mg/mL chitosan in 0.1 M acetic acid.

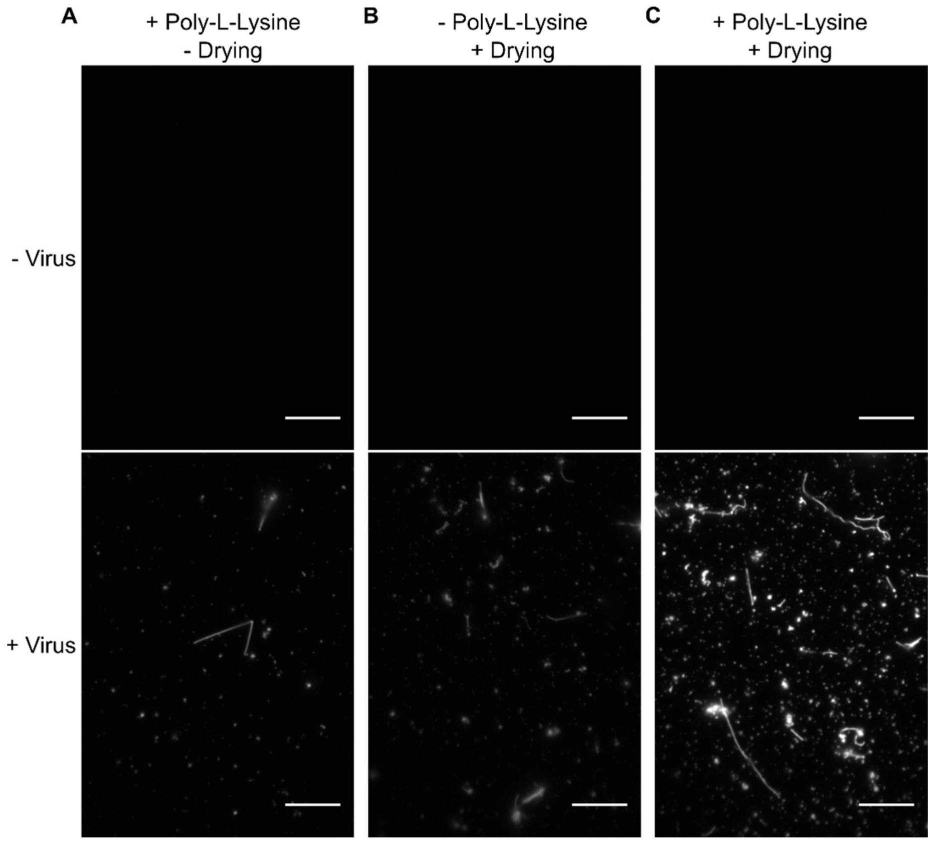


**Sup. Fig. 2. A combination of drying the sample and using poly-L-lysine increases the number of immobilized viruses on the slide surface.** A) A virus negative control (top) or an A/Udorn/72 virus sample (bottom), were incubated on a slide pre-treated with 0.01% poly-L-lysine at 4°C for 10 minutes. The excess sample was removed from the well and the immobilized virus was fixed and labelled with an anti-Udorn primary antibody and an Alexa647 secondary antibody before being imaged on a widefield TIRF microscope. Scale bars 10µm. B) As in A) but the slide was not pre-treated with poly-L-lysine and the samples were dried directly onto glass coverslips by heating at 45°C for 10 minutes. C) As in A) but the samples were dried directly onto glass coverslips by heating at 45°C for 10 minutes.

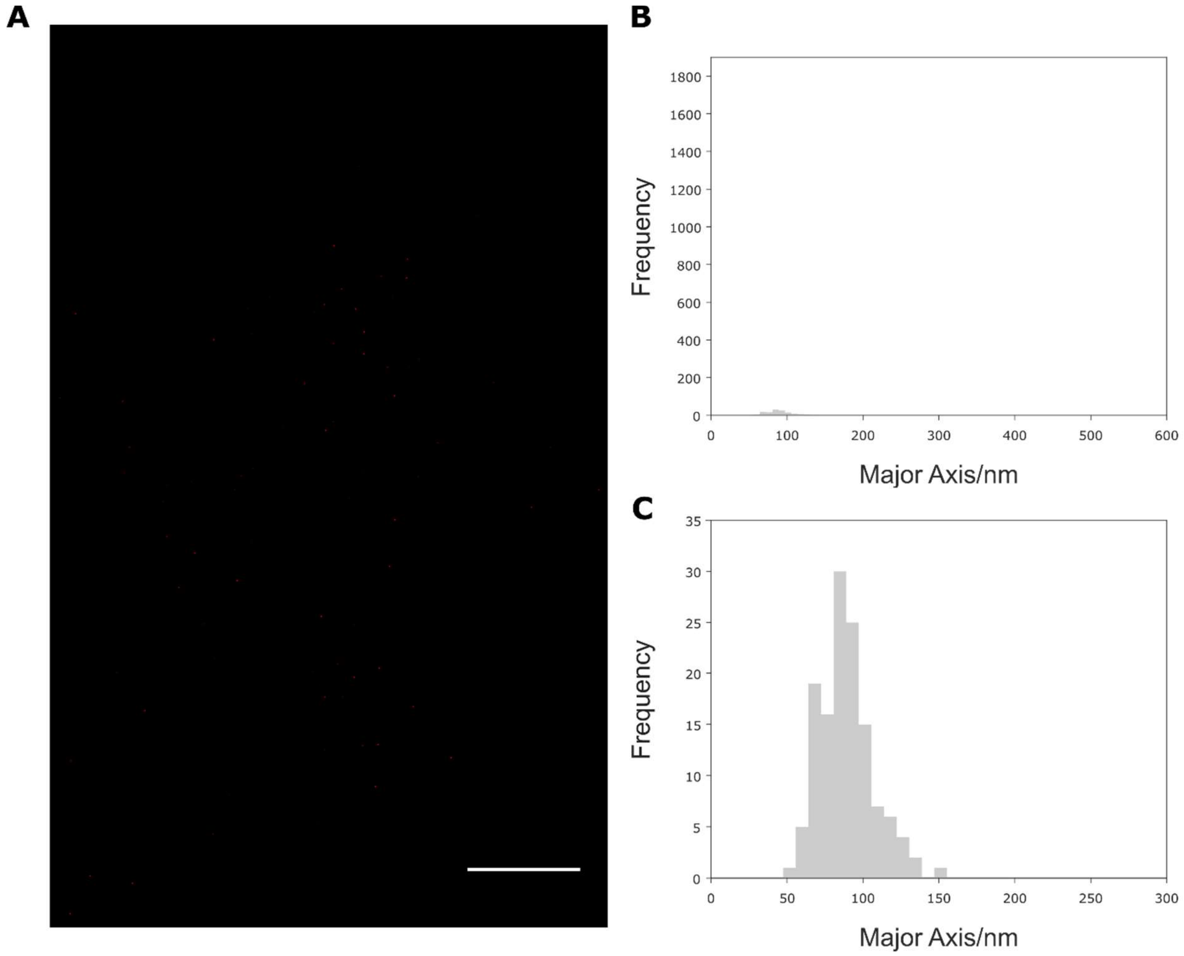


**Sup. Fig. 3. Virus-negative FOV and histograms for the size analysis of spherical and bacilliform influenza particles.** A) A representative super-resolution image of a virus-negative sample virus stained with an antibody against the HA protein. Scale bar 10 µm. B) Super-resolution localisations were clustered and each cluster fitted with an ellipse to extract particle dimensions. A histogram of the major axis lengths shows that background signal is negligible. C) Zoomed in histogram of B).

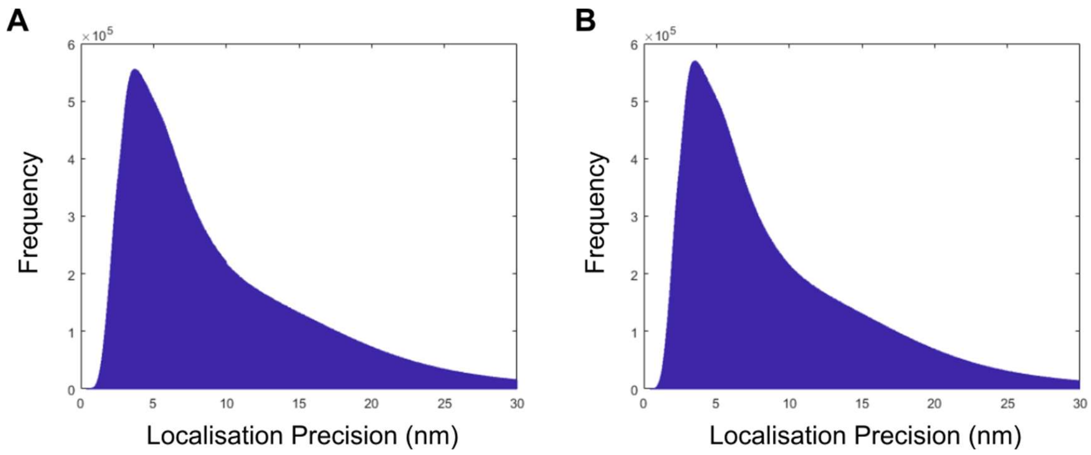


**Sup. Fig. 4. Histograms of the localisation precisions of the data from Figure 3.** A) Each FOV was drift corrected, and a Gaussian function was fitted to each detected localization in every frame of the acquisition, using the inbuilt Nanoimager software. The fitting error (or localization precision) in the x direction of each localization was exported and plotted as a histogram, providing a median error of 7.4 nm. B) Plot of the localisation precision in the y direction, providing a median error of 7.2 nm.

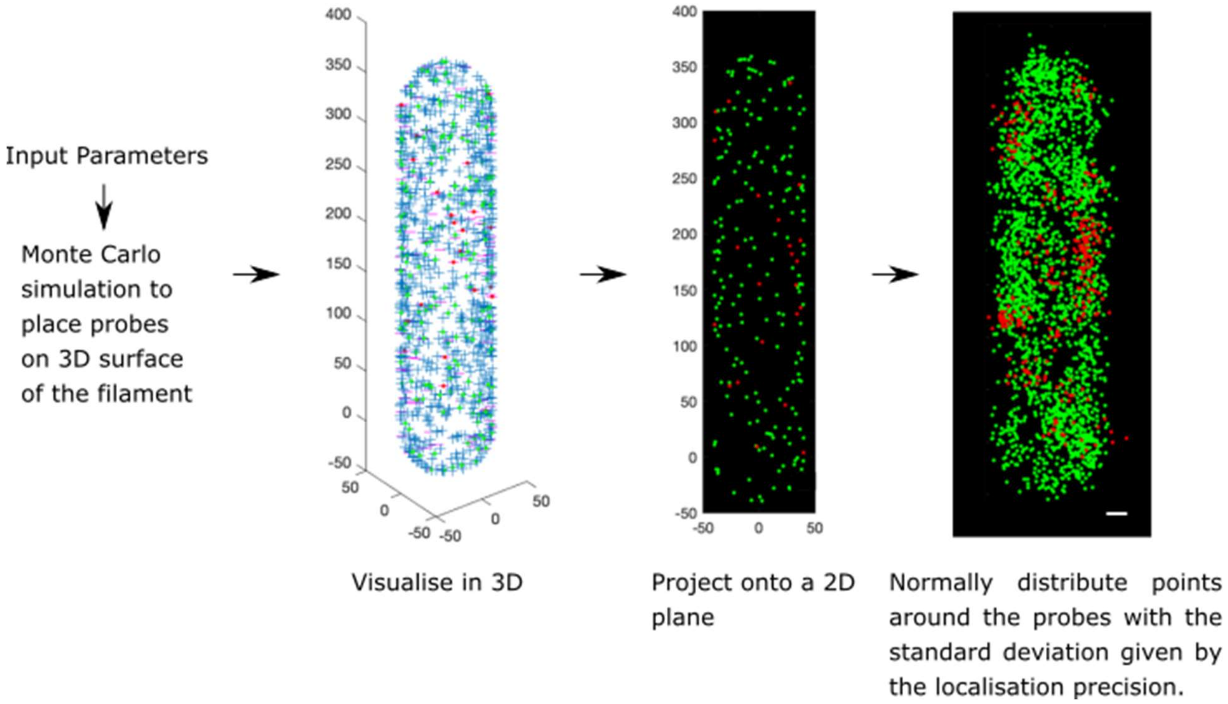


**Sup. Fig. 5. Summary of the dSTORM simulation method.** Input parameters of virion size, number of localisations and localization precision were used for Monte Carlo simulations to create simulated dSTORM images of filamentous virions. Filaments were modelled as cylinders with hemispherical caps. After projecting the simulated filaments into 2D, localisations were randomly placed with a normal distribution about the protein location with a standard deviation of 7.4nm and the image was coloured as a STORM image. Scale bar 20nm.

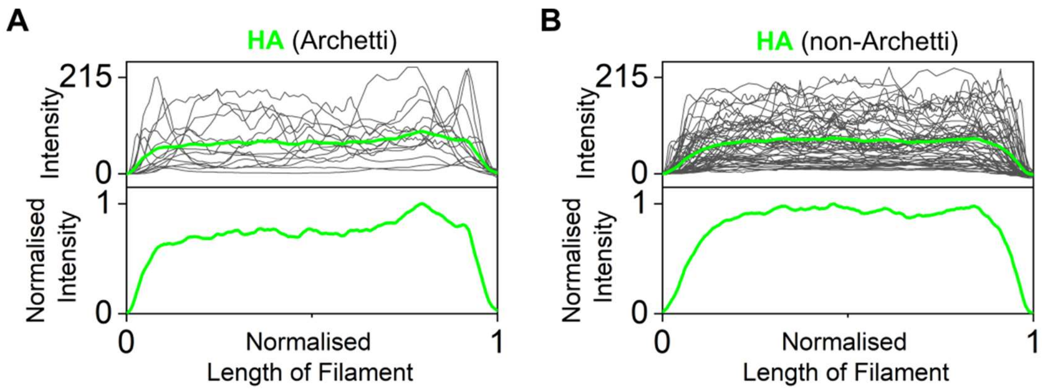


**Sup. Fig. 6. HA distribution of filaments shows when Archetti bodies are present.** A) Top: Intensity traces (grey) of the HA signal from 14 filaments with a visible Archetti body, with the average intensity profile shown as a green line. Bottom: Normalised average RNA signal from the 14 filaments. B) Same as A) but for the HA signal from 54 filaments with no visible Archetti body.
